# Comparative genomic analyses provide clues to capsule switch in *Streptococcus suis*

**DOI:** 10.1101/2020.11.11.377622

**Authors:** Yinchu Zhu, Wenyang Dong, Jiale Ma, Yue Zhang, Xiaojun Zhong, Zihao Pan, Guangjin Liu, Zongfu Wu, Huochun Yao

**Affiliations:** Institute of Animal Husbandry and Veterinary Sciences, Zhejiang Academy of Agricultural Sciences, Hangzhou 310021, China; College of Veterinary Medicine, Nanjing Agricultural University, Nanjing 210095, China; Key Lab of Animal Bacteriology, Ministry of Agriculture, Nanjing 210095, China; OIE Reference Lab for Swine Streptococcosis, Nanjing 210095, China; Beijing Advanced Innovation Center for Genomics (ICG) & Biomedical Pioneering Innovation Center (BIOPIC), Peking University, Beijing 100871, China; College of Animal Science and Veterinary Medicine, Henan Agricultural University, Zhengzhou 450002, China; College of Animal Science and Technology, College of Veterinary Medicine, Zhejiang A & F University, Hangzhou, 311300, China

**Keywords:** Streptococcus suis, Capsule switch, S. suis serotype 2, S. suis serotype 3, S. suis serotype 7, MLST, Comparative genomic analyses

## Abstract

*Streptococcus suis* (*S. suis*) is a major bacterial pathogen in swine industry and also an emerging zoonotic agent. *S. suis* produces an important extracellular component, capsular polysaccharides (CPS). Based on which, dozens of serotypes have been identified. Through virulence genotyping, we uncovered the relatedness between proportions of SS2, SS3 and SS7 strains despite their differences in serotypes. Multi-locus sequence typing (MLST) was used to characterize whole *S. suis* population, revealing that there is capsule switch between *S. suis* strains. Importantly, capsule switch occurred in SS2, 3 and 7 strains belonging to CC28 and CC29, which is phylogenetically distinct from the main CC1 SS2 lineage. To further explore capsule switch in *S. suis*, comparative genomic analyses were performed using available *S. suis* complete genomes. Phylogenetic analyses suggested that SS2 strains can be divided into two clades (1 and 2), and those classified into clade 2 are colocalized with SS3 and SS7 strains, which is in accordance with above virulence genotyping and MLST analyses. Clade 2 SS2 strains presented high genetic similarity with SS3 and SS7 and shared common competence and defensive elements, but are significantly different from Clade 1 SS2 strains. Notably, although the *cps* locus shared by Clade 1 and 2 SS2 strains is almost the same, a specific region in *cps* locus of strain NSUI002 (Clade 2 SS2) can be found in SS3 *cps* locus, but not in Clade 1 SS2 strain. These data indicated that SS2 strains appeared in CC28 and CC29 might acquire *cps* locus through capsule switch, which could well explain the distinction of genetic lineages within SS2 population.

## 1. Introduction

*Streptococcus suis* (*S. suis*) is a major bacterial pathogen causing global economic losses to swine industry. It is also a serious zoonotic pathogen in countries with intensive swine production. Capsule polysaccharide (CPS) is the key virulence determinant in *S. suis*, which contributes to the bacterial resistance to host immunity (Fittipaldi et al., 2012). Sequence analyses of *S. suis* genomes revealed a *cps* locus with variable lengths and a series of genes specific to CPS production (Okura et al., 2013). It is thought that CPS of *S. suis* is synthesized and exported through the Wzx/Wzy pathway, which is a mechanism commonly used in *Streptococcus pneumoniae* (*S. pneumoniae*) and *Streptococcus agalactiae* (*S. agalactiae*) capsular biosynthesis (Yother, 2011). The difference in *cps* locus would lead to the difference in components and structures of CPS, and importantly, the serum antigenicity. In the 1980s and 1990s, 35 serotypes (types 1 to 34 and type 1/2) having been described based on CPS antisera coagglutination test (Higgins et al., 1995). Later on, serotypes 20, 22, 26, 32, 33 and 34 are suggested to be removed from *S. suis* species (Hill et al., 2005; Tien et al., 2013). More recently, a novel variant serotype Chz and other 8 novel *cps* loci harboring specific *wzy* polymerase genes and *wzx* flippase genes were identified (Pan et al., 2015; Qiu et al., 2016), revealing the high diversity of *cps* locus in *S. suis* genomes.

Among the known serotypes worldwide, *S. suis* serotypes 2, 3, 9, 7, 8, 4 and 1 are the most prevalent serotypes linked with infection of swine, especially serotype 2 (SS2) (Goyette-Desjardins et al., 2014). In North America, serotype 2 and 3 are the two most prevalent serotypes isolated from clinical pig cases (Goyette-Desjardins et al., 2014). In Asia, the most prevalent serotypes in infected pigs are serotypes 2, 3, 4, 7 and 8 (Goyette-Desjardins et al., 2014; Wei et al., 2009). In Europe, serotype 9 and 2 are more frequently found in clinical pig cases, followed by serotypes 7, 8, 3 and 1 (Goyette-Desjardins et al., 2014). As for the infection of human, serotypes 2, 4, 5, 9, 14, 16, 21 and 24 have already been reported, of which SS2 is the predominate serotype (Goyette-Desjardins et al., 2014; Kerdsin et al., 2017). Therefore, most studies in *S. suis* field are focusing on SS2 due to its close link with diseases. However, heterogenicity in SS2 population has been observed, indicating serotyping alone is not sufficient to characterize *S. suis* strains.

In addition to capsule serotyping, multilocus sequence typing (MLST) is most the widely used typing method in epidemiological studies of *S. suis*. Analyses of MLST data from different sources determined the sequence types (STs) of stains, which could be further clustered into clonal complexes (CCs) (King et al., 2002). Interestingly, MLST analyses of *S. suis* isolates suggested that SS2 population can be divided into two major lineages, not only in terms of geographical and genetic background, but also virulence phenotype (Fittipaldi et al., 2011; Goyette-Desjardins et al., 2014; Yao et al., 2015; Zhu et al., 2013). Among SS2 isolates, although ST1/ST7 (CC1), ST25 (CC29) and ST28 (CC28) strains have been isolated in both Asia and North American, CC1 strains are more prevalent in Asia, while CC28 and CC29 strains are more commonly found in North America (Goyette-Desjardins et al., 2014). Importantly, ST1/ST7 strains are significantly more virulent than ST25 and ST28 strains (Athey et al., 2015; Fittipaldi et al., 2011; Guo et al., 2020). Although the distinctions of genetic and virulence phenotypic lineages within SS2 population have been found, the way it was formed are not yet fully understood.

Capsule switch, a change of serotype of a single clone by alteration or exchange of its *cps* locus, has been identified in streptococcal species including *Streptococcus iniae* (*S. iniae*) (Heath et al., 2016), *S. pneumoniae* (Wyres et al., 2013) and *S. agalactiae* (Martins et al., 2010). Development of MLST greatly promote the studies on capsular switch, which can be more easily identified by detecting strains of different serotypes sharing the same ST. Although capsular switch has been found in *S. suis* isolates (King et al., 2002), the effects of this phenomena on *S. suis* population structure has not been demonstrated. Previously, we showed that the virulence related genes are differently distributed in strains of two SS2 clusters (Dong et al., 2015), which is in accordance with the known SS2 sequence type classification (Fittipaldi et al., 2011; Zhu et al., 2013). In this study, we combined MLST analysis, virulence genotyping and whole genome analysis, to explore the capsular switch in *S. suis*, and highlight its potential role of in shaping SS2 population.

## 2. Materials and methods

### 2.1 Bacteria strains and culture conditions

All *S. suis* serotype 3 (SS3) and *S. suis* serotype 7 (SS7) strains are the field strains isolated from China from 2004 to 2018 and were stored in our laboratory. The typical virulent SS2 strain ZY0719 was isolated from a diseased pig during an outbreak in China. *S. suis* was grown in Todd–Hewitt broth medium (THB; Becton Dickinson, Sparks, MD, USA) at 37°C overnight. The antibiotics including spectinomycin (100□μg/ml) and chloramphenicol (5 μg/ml) were added into the medium if it is needed. The plasmids for Streptococcus pSET-2::cat was used in this experiment for DNA template. A detailed information for bacterial strains used in this study listed in Table S1.

### 2.2 PCR assays

A previously established species-specific polymerase chain reaction (PCR) based on the *gdh* and *recN* genes were performed to confirm the identification of *S. suis* (Okwumabua et al., 2003). The serotype-specific PCR was used to identify SS3 and SS7 strains among the collected *S. suis* isolates (Kerdsin et al., 2014). In the virulence genotyping assay, 19 *S. suis* virulence-associated genes, including *mrp*, *epf, sly, rgg, ofs, srtA, pgdA, gapdh, iga, endoD, ciaRH, salKR, manN, purD, dppIV, neuB, dltA, comR* and *scnF* were detected by individual PCR as previously described (Dong et al., 2015; Dong et al., 2017). In MLST assays, seven housekeeping genes *aroA, mutS, cpn60, dpr, recA, thrA* and *gki* were amplified by PCR as described previously (King et al., 2002), and the amplification fragments were sequenced.

### 2.3 Clustering analysis

For MLST, allele numbers and sequence types (STs) were identified in MLST database (http://ssuis.mlst.net/). The eBURST (http://eburst.mlst.net) program was used to determine population structures through identifying potential clonal complexes (CCs) and founders. For virulence genotyping, BioNumerics (version 6.6, Applied Maths, Kortrijk, Belgium) was used to analyze the profiles of virulence related genes as previously described (Dong et al., 2015; Dong et al., 2017; Mateus et al., 2013; Zhu et al., 2017): the resemblance was computed with simple matching coefficients, and agglomerative clustering was performed using the unweighted average linkage (UPGMA). The profiling of SS2 virulence related genes used in this clustering analysis is acquired from a previous study (Dong et al., 2015).

### 2.4 Phylogenetic analysis

The complete genome sequences of 25 *S. suis* isolates of different serotypes were obtained from NCBI GenBank and used for phylogenetic analyses, including the 3 *S. suis* genomes provided by our research group (ZY05719, T15 and SC070731). A phylogenetic tree based on the 1373 single copy orthology clusters generated from clustering of the 25 strains was constructed using neighbor-Joining (NJ) method. The information of strains were listed in table S2.

### 2.5 Multi-genome alignment analysis

Comparative analysis of alignments among *S. suis* genomes were conducted using the progressive alignment option of the Mauve software (Darling et al., 2004). ZY05719 was selected as virulent SS2 strains, NSUI002 and NSUI060 were the typical avirulent SS2 strains. Genomes of SS3 strains YB51 and ST3, SS7 strain D9 were also used in genome alignments.

### 2.6 Average nucleotide identity analysis

The genomic similarity between SS2, SS3 and SS7 strains was evaluated by average nucleotide identity (ANI) method. The algorithm implemented at the EzGenome server was used To calculate the ANI value (www.ezbiocloud.net/tools/ani). The proposed and generally accepted species boundary for ANI value are 95~96% (Richter and Rossello-Mora, 2009).

### 2.7 Defense elements analysis

Bacterial defense systems were ancient elements that confers resistance to foreign genetic elements, including Restriction-Modification (RM) system and Clustered Regularly Interspaced Short Palindromic Repeats (CRISPR) system. RM elements and CRISPR components in *S. suis* were determined through CRISPRs finder (http://crispr.u-psud.fr/) and web-service REBASE (http://rebase.neb.com/rebase/rebase.html), representatively. The structures of RM and CRISPR systems are achieved through performing an all-to-all BLASTN search in the NCBI nucleotide database. Consensus sequences of repeat sequences in defense systems were determined using MegAlign.

### 2.8 Competence system analysis

*S. suis* is harbors a natural competence system, which greatly facilitates the DNA exchange through horizontal genetic transfer. In our previous study, we identified that ComRS-ComX competence system varies in different genotype background *S. suis* strains (Zhu et al., 2019). Through BLASTN search, competence systems were identified in representative SS2, SS3, SS7 genomes. The amino acid sequence of competence systems was visualized by DNAMAN software. To experimentally validate the functional difference between *S. suis* strains, a transformation efficiency test was performed between 8 selected SS2, SS3 and SS7 field strains stored in our laboratory. Briefly, two different synthetic competence peptides XIP (type A:GNWGTWVEE and type B: LGDENWWVK) were added to the bacterial culture with template DNA (plasmid pSET-2::cat) to induce the natural transformation. Competency was calculated based on the amount of transformants that grew in THB plates with spectinomycin and chloramphenicol selection.

### 2.9 Analysis of *cps* locus

The diversity serotypes are based on the variation of genetic locus harboring capsular polysaccharide related genes (*cps* locus). In general, strains with a same serotype would present highly similar genomic sequence in *cps* locus. The sequence of *cps* locus in genomes of CC1 SS2 strain ZY05719, CC28 SS2 strain NSUI002, and CC28 SS3 strain ST3 were determined using BLAST. Corresponding sequences and gene annotation information were obtained from obtained from NCBI GenBank. The homologous analysis was performed using Easyfig software.

## 3. Results

### 3.1 Virulence genotyping of field strains revealed relatedness of a SS2 subpopulation with SS3 and SS7

Virulence genotyping is a powerful tool to study pathogenic bacteria, which can contribute to screen for specific disease-associated virulence genes (Gerjets et al., 2011; Rasmussen et al., 2013) or uncover the relatedness between isolates (Dong et al., 2017; Mateus et al., 2013). Previously, we applied a virulence genotyping strategy by detecting a set of virulence related genes in SS2 isolates, and revealed that SS2 strains can be divided into two clusters due to the different distribution of genes *epf, sly, endoD, rgg* and *scnF* (Dong et al., 2015). In this study, we further detected the presence of virulence related genes in SS3 and SS7 isolates by PCR, and performed clustering analysis based on the gene profiles of SS2 (n=62), SS3 (n=17) and SS7 (n=9) isolates (Figure 1). In accordance with what we have showed previously (Dong et al., 2015), SS2 strains could be divided into two clusters ((I and II) with a different prevalence of virulence related genes. Interestingly, SS3 and SS7 isolates were classified into cluster II together with one SS2 subpopulation, suggesting a relatedness of SS3 and SS7 with that SS2 subpopulation (Figure 1). For the gene distribution, 8 genes were detected in all *S. suis* field isolates, namely *srtA, pgdA, dltA, iga, sspA, manN, ciaHR* and *gapdh*; whereas gene *epf* and *rgg* were only detected in cluster I but not in cluster II. According to virulence genotyping result in this study and previous knowledge (Dong et al., 2015; Dong et al., 2017; Kobayashi et al., 2013; Mateus et al., 2013), we hypothesized that the presence of similar virulence genotypes reflects phylogenetic relatedness of SS2, SS3 and SS7. Therefore, we performed genetic analyses in *S. suis* population to further test our hypothesis.

**Figure 1.**
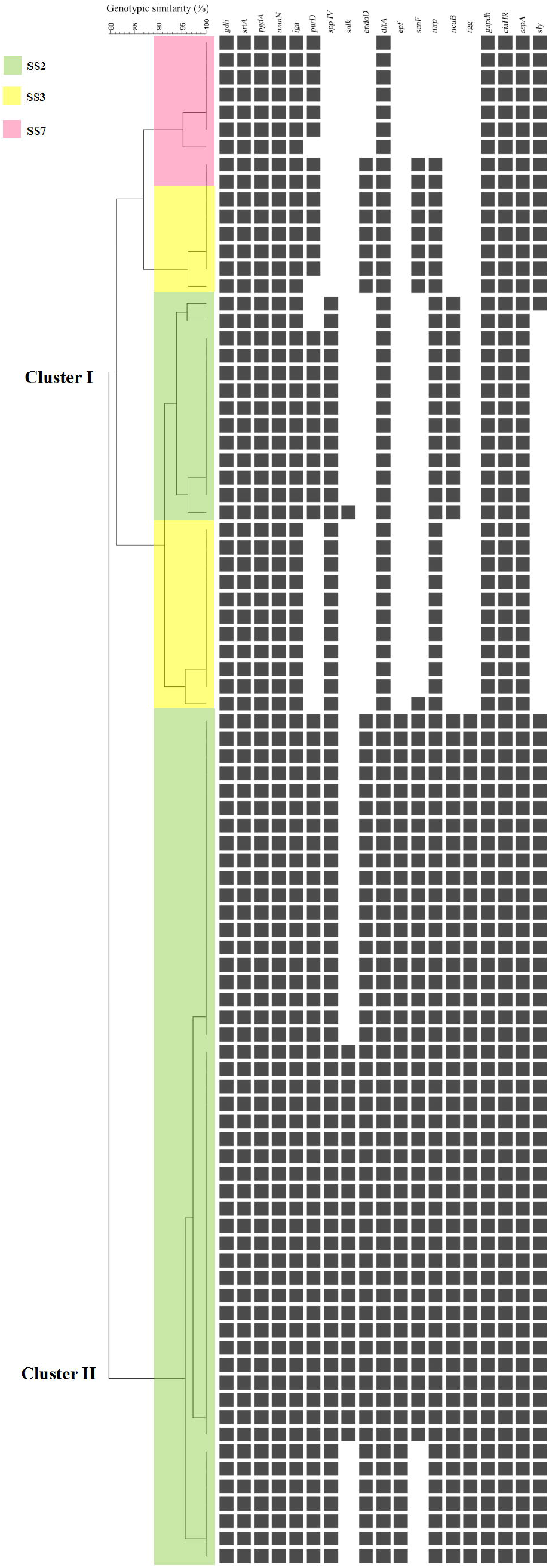
Clustering of SS2, SS3 and SS7 isolates based on the profiles of virulence related genes. The lateral axis of the matrix is the set of 19 genes and the vertical axis is the set of 88 strains. Green color covers SS2 strains, yellow covers SS3 strains and pink covers SS7 strains. Each black square refers to a positive detection of a specific virulence related gene in a single *S. suis* strain. Agglomerative clustering was performed using the unweighted average linkage (UPGMA) with the BioNumerics software.

### 3.2 MLST analysis demonstrated capsule switch in *S. suis* between SS2, SS3 and SS7

MLST is an important and widely used molecular method in studying *S. suis* epidemiology, in which seven housekeeping genes are sequenced to assess the genomic variation and define Sequence Types (STs). The availability of updated information in *S. suis* MLST database (http://ssuis.mlst.net/) makes it possible to apply a grouping approach for identification of Clonal Complex (CC) and perform comparative analysis with strains of different serotypes. A total of 1528 *S. suis* strains (701 STs) from MLST database was diagramed by eBURST on the basis of their allelic profiles. The eBURST analysis revealed 9 major clonal complexes (Figure 2A). CC1 (629/1528) is the predominant clonal complexes in *S. suis*, followed by CC16 (120/1528), CC28 (71/1528), CC29 (62/1528) and CC104 (30/1528), whereas CC94, CC528, CC423 and CC201 are much smaller schemes.

**Figure 2.**
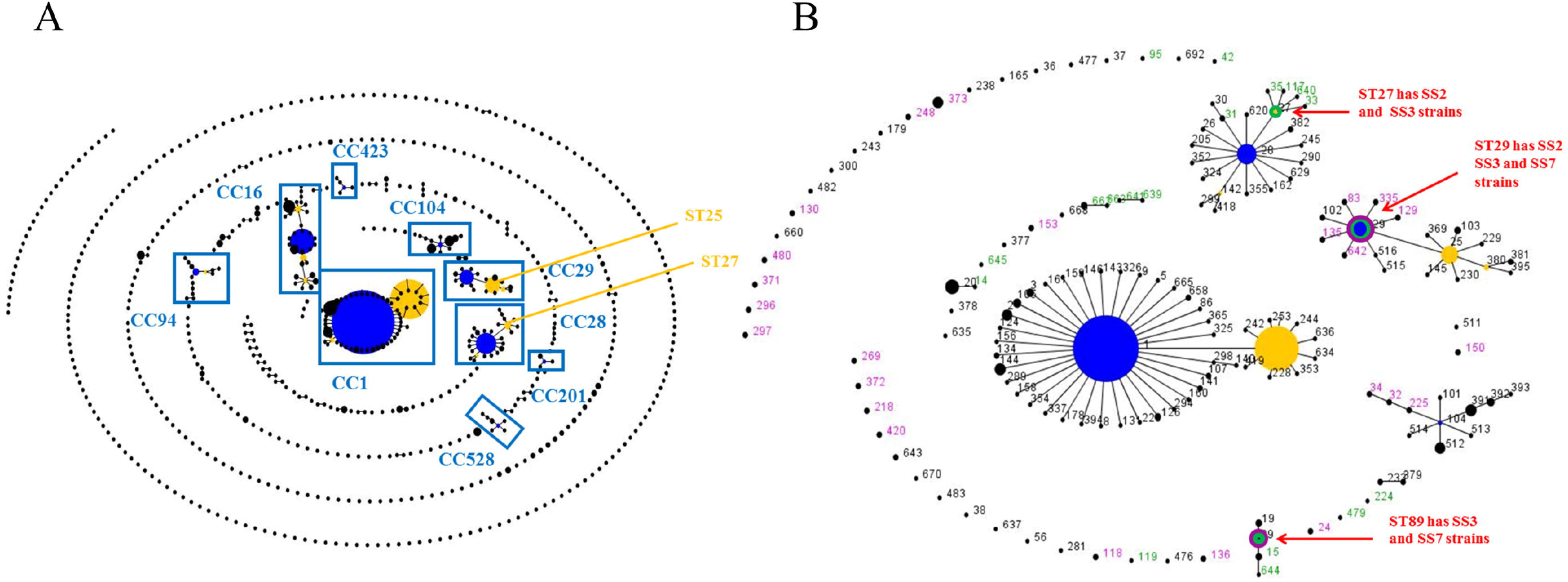
eBURST analysis of *S. suis* MLST data. **(A)** eBURST software was used to analyze the MLST data of whole *S. suis* population. A total of 9 major CCs were identified, including CC1, CC16, CC28, CC29, CC94, CC104, CC201, CC423 and CC528. **(B)** Detailed analysis all SS2, SS3 and SS7 strains. SS2 strains are colored in back, SS3 in green and SS7 in purple. ST1 and ST7 SS2 are clustered into CC1. CC28 includes both SS2 and SS3 strains, whereas CC29 contains SS2 and SS7 strains.

The MLST analysis revealed capsular switch in *S. suis*, that strains of different serotypes sharing the same ST. In *S. suis*, capsular switch was firstly reported in 2002 (King et al., 2002), which analyzed the capsular switch events from ST1 to ST92, and showed it occurred in ST1, 13, 16, 17, 27, 28, 29, 65 and 76. Our updated analysis identified novel STs with capsular switch, including ST15, 89, 94, 105, 136, 156, 243 and 297 (Table 1). Importantly, there is capsular switch within SS2, SS3 and SS7 strains. ST27 from CC28 has both SS2 and SS3 strains, and ST29 from CC29 has SS2, SS3 and SS7 strains (Table 1 and Figure 2B). CC28 and CC29 have already been demonstrated to be important SS2 lineages (Athey et al., 2015; Fittipaldi et al., 2011). Interestingly, as shown in our eBURST analysis, CC28 harbors a SS3 major population, and CC29 harbors a SS7 major population (Figure 2B). Those results suggested that a sub-population of SS2 strains of CC28 and CC29 are more closely related to SS3 and SS7 strains, and there is capsular switch in those two clonal complexes.

**Table 1.**
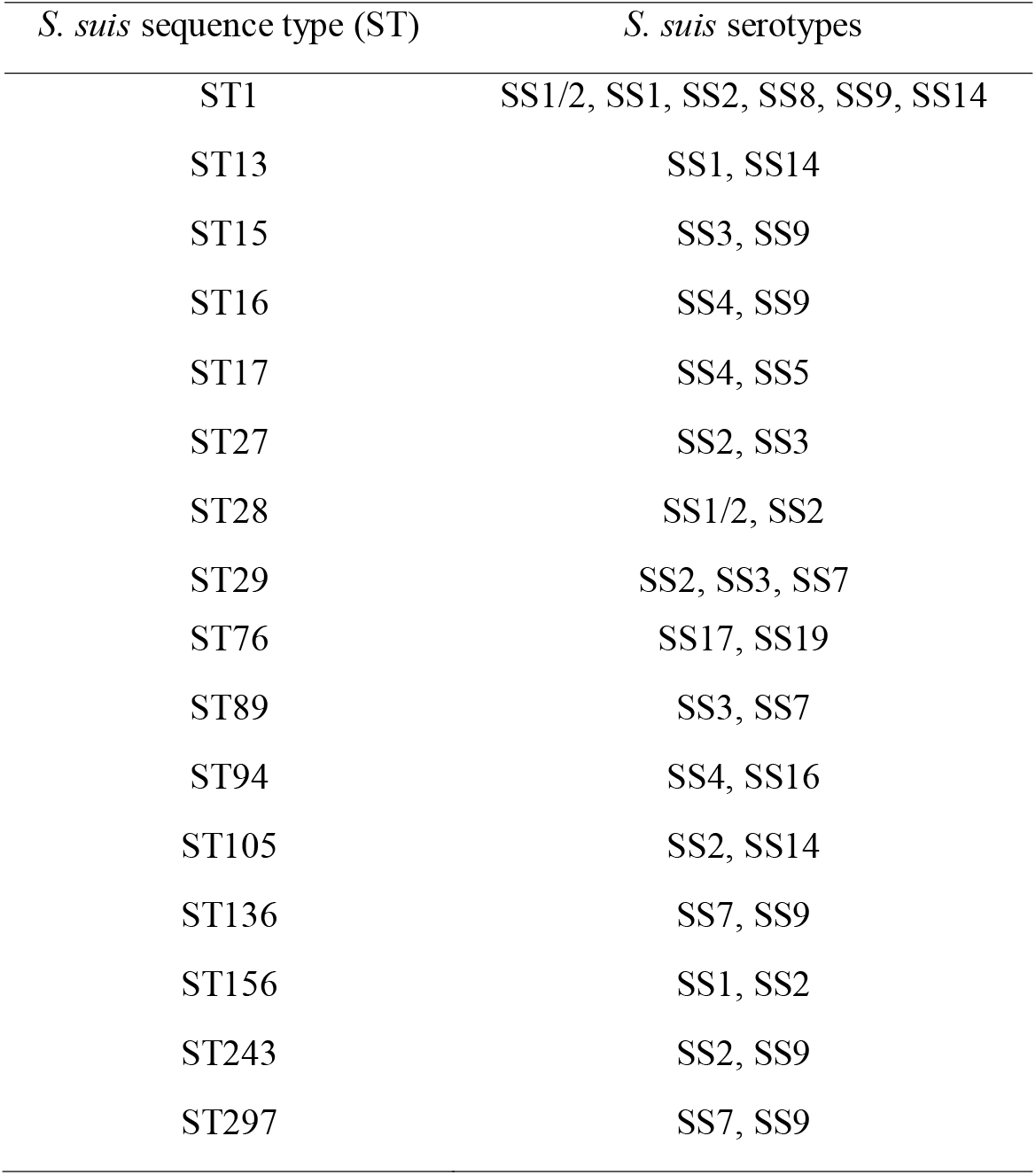
Existing capsule switch in *S. suis*

### 3.3 Whole genome phylogenetic analysis of different serotype strains deciphered two distinct clades in SS2

To investigate the phylogenetic relationships among *S.suis* strains, we used 25 complete genomes of different serotypes from GenBank dataset. The phylogenetic tree, constructed using neighbor-joining (NJ) method (Figure 3), showed that *S.suis* strains of different serotypes can be classified into three major clades (Clade 1, 2 and 3). SS1 strain, SS4 strain and SS2 avirulent strain T15 appeared to be phylogenetically independent, whereas SS16 strain and SS9 strains are located together in Clade 3.

**Figure 3.**
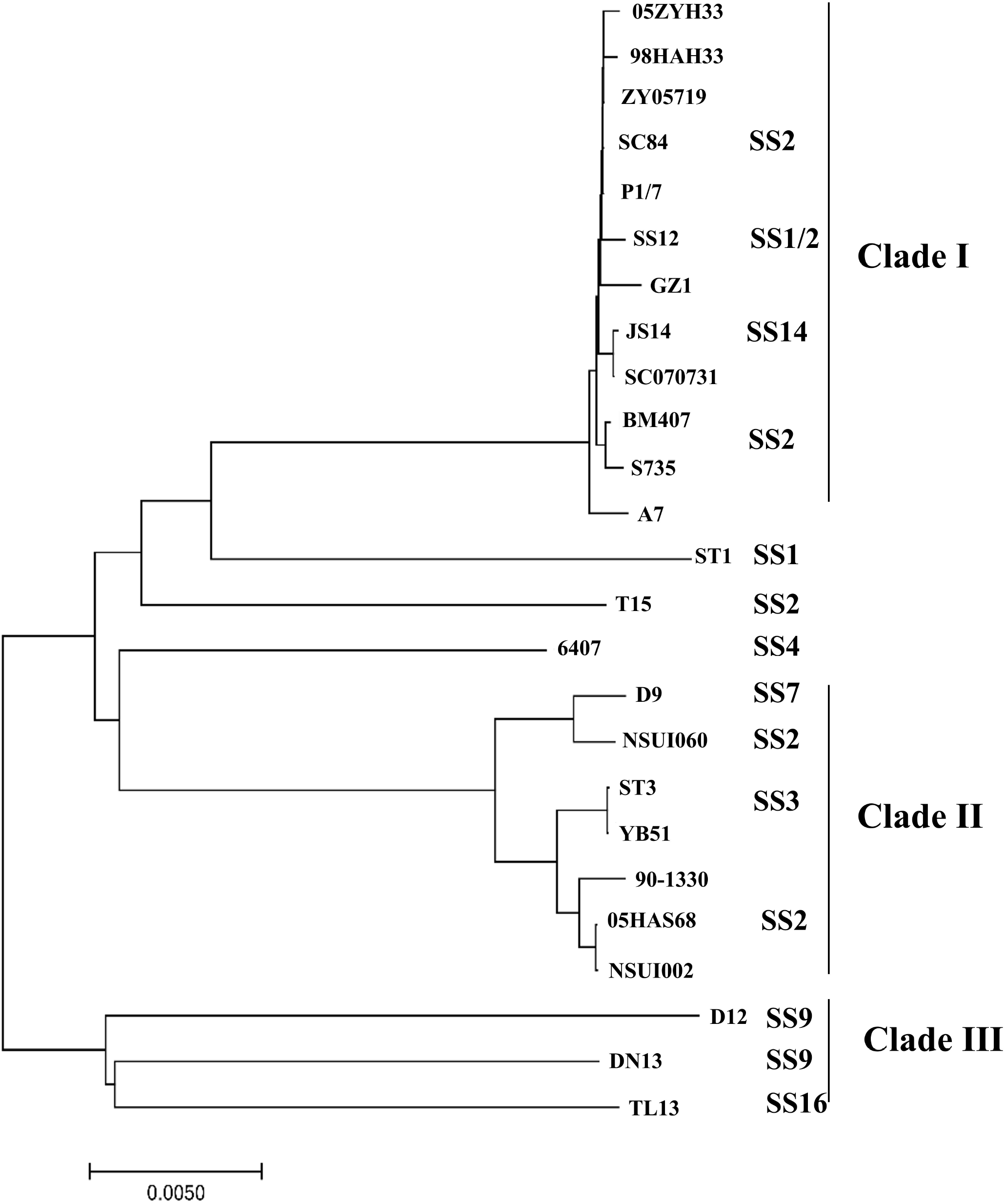
Phylogenetic tree of 25 *S. suis* strains based on orthologous gene clusters. The phylogenetic tree was constructed using neighbor-joining (NJ) method. *S. suis* strains of different serotypes can be classified into 3 major clades. Clade 1 appeared to be phylogenetic distinct from Clade 2. Clade 1 harbors virulent SS2 strains, and Clade 2 SS3 strains, SS7 strain and SS2 strains with lower virulence level.

SS14 strain JS14 and SS1/2 strain SS12 are grouped together with ten SS2 strains, and formed the largest clade (Clade 1) on phylogenetic tree. SS2 strains in this clade presented a short evolutionary distance from each other, suggesting that these strains were probably derived from a recent common ancestor. This result is in accordance with the MLST analysis, that all SS2 strains in Clade 1 belong to CC1. SS14 strain JS14 and SS1/2 strain SS12 also presented CC1 related STs, indicating a close link with Clade 1 SS2 strains.

Clade 2 included SS7 strain D9, SS3 strains ST3 and YB51, and four SS2 strains. These SS2 strains therefore showed a large divergence from Clade1 SS2. SS2 strain NSUI060 is a strain of ST25 (CC29), and is assigned in a same branch with ST29 (CC29) strain D9 on phylogenetic tree. SS2 strains 90-1330, 05HAS68 and NSUI002 belongs to ST28 (CC28), and are grouped in a same branch with two ST 35 (CC28) SS3 strains. Above result indicated that CC28/29 SS2 strains in this clade were more closely related with SS7 strain and SS3 strains rather than CC1 SS2 strains in Clade 1. This is consistent with the virulence genotyping result, and also reflecting the capsule switch identified by MLST within SS2, SS3 and SS7strains. Therefore, we further performed whole-genome comparative analysis to decipher capsule switch in *S. suis*.

### 3.4 Clade 2 SS2 had a higher genomic similarity with SS3 and SS7 than Clade 1 SS2

The arrangement and collinearity of Clade 1 and Clade 2 *S. suis* genomes were investigated using Mauve program. We first used CC28 SS2 strain NSUI002 as reference genome, to compare clade 1 CC1 SS2 strain ZY05719, as well as CC28 SS3 strains ST3 and YB51 that colocalized with NSUI002 in Clade 2 sub-branch. Mauve analysis (Figure 4) of those strains indicated that rearrangements occurred in genomes of those strains, but the overall genomic organizations are relatively comparable. The NSUI002 and YB51 genomes showed more collinear than ZY05719, suggesting that NSUI002 and SS3 strains have a higher genomic similarity. We further compared the genomic organization of SS2 strain NSUI060 and SS7 strain D9 (Figure S1), which are colocalized in a subgroup of Clade 2 as well. Alignment of the genome sequences of NSUI060 and D9 revealed a high level of genomic rearrangement, including large-scale deletion, insertion, translocation, and inversion. However, the genomes of NSUI060 and D9 still shared a more similar sequence synteny with each other, but significantly different from ZY05719 with respect to both collinearity and genome structure. Those results show that Clade 2 SS2 strains and SS3/7 strains has the smaller scale of arrangements and higher level of synteny compared with Clade 1 SS2.

**Figure 4.**
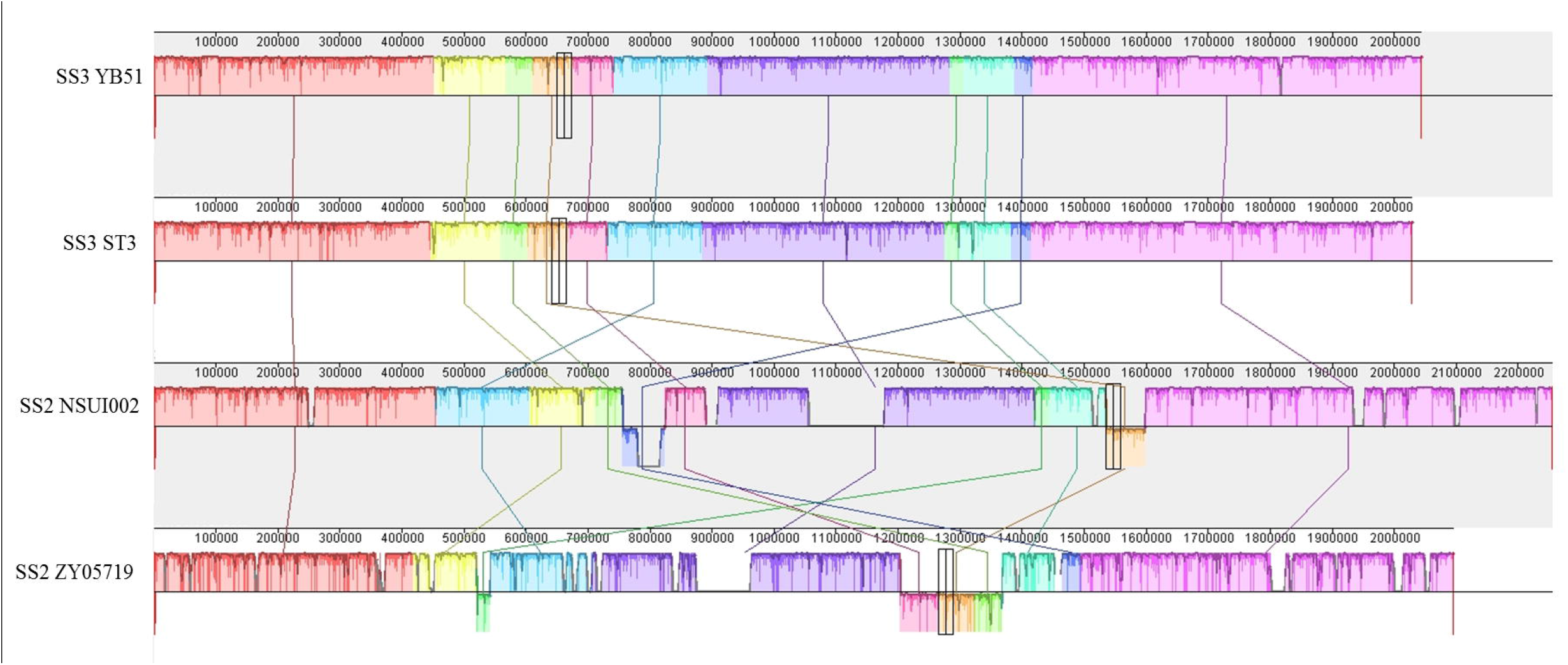
Multigenome comparison between Clade 1 and Clade 2 *S. suis* strains obtained by Mauve tool. Each colored region refers to a locally collinear block (LCB). Colors are arbitrarily assigned by software to each LCB. The vertical peaks in each LCB denotes the variance of conservation. The LCBs below the center line of genomes are in reverse complement orientation. As reference genome, Clade 2 SS2 strain NSUI002 are compared with SS3 strains (ST3 and YB51) and Clade 1 SS2 strain ZY05719. The Multigenome comparison of SS2 strain ZY05719, NSUI060 and SS7 strain D9 is shown in Figure S1.

The genomic relationship between Clade 1 isolates and Clade 2 isolates was further evaluated by nucleotide sequence similarity. The average nucleotide identity (ANI) were calculated (Figure 5) using the web-based EZ BioCloud platform (www.ezbiocloud.net/tools/ani). The ANI values between the Clade 1 SS2 isolates are very high, ranged from 99.54 to 99.97□%, whereas the ANI values between Clade 1 SS2 isolates and Clade 2 SS2 is only from 96.28 to 96.94, which is close to the cut-off values recommended for species delineation (95–96□≥%) (Richter and Rossello-Mora, 2009). The Clade 2 SS2, SS3 and SS7 strains have ANI values above 98, and can be further divided into two sub-clusters, which is in accordance with the phylogenetic analysis and Mauve result. Especially, the ANI values between three Clade 2 SS2 isolates, namely 90-1330, 05HAS68 and NSUI002, and SS3 strains are from 99.57 to 99.70, while the ANI value between SS2 strain NSUI060 and SS7 strain D9 is 99.50. Thus, these results revealed the genomic dissimilarity of Clade 1 SS2 isolates and Clade 2 SS2 isolates, supporting the conclusion that Clade 2 SS2 isolates have a high relatedness with SS3 and SS7 strains, as implied by the above whole genome comparison results.

**Figure 5.**
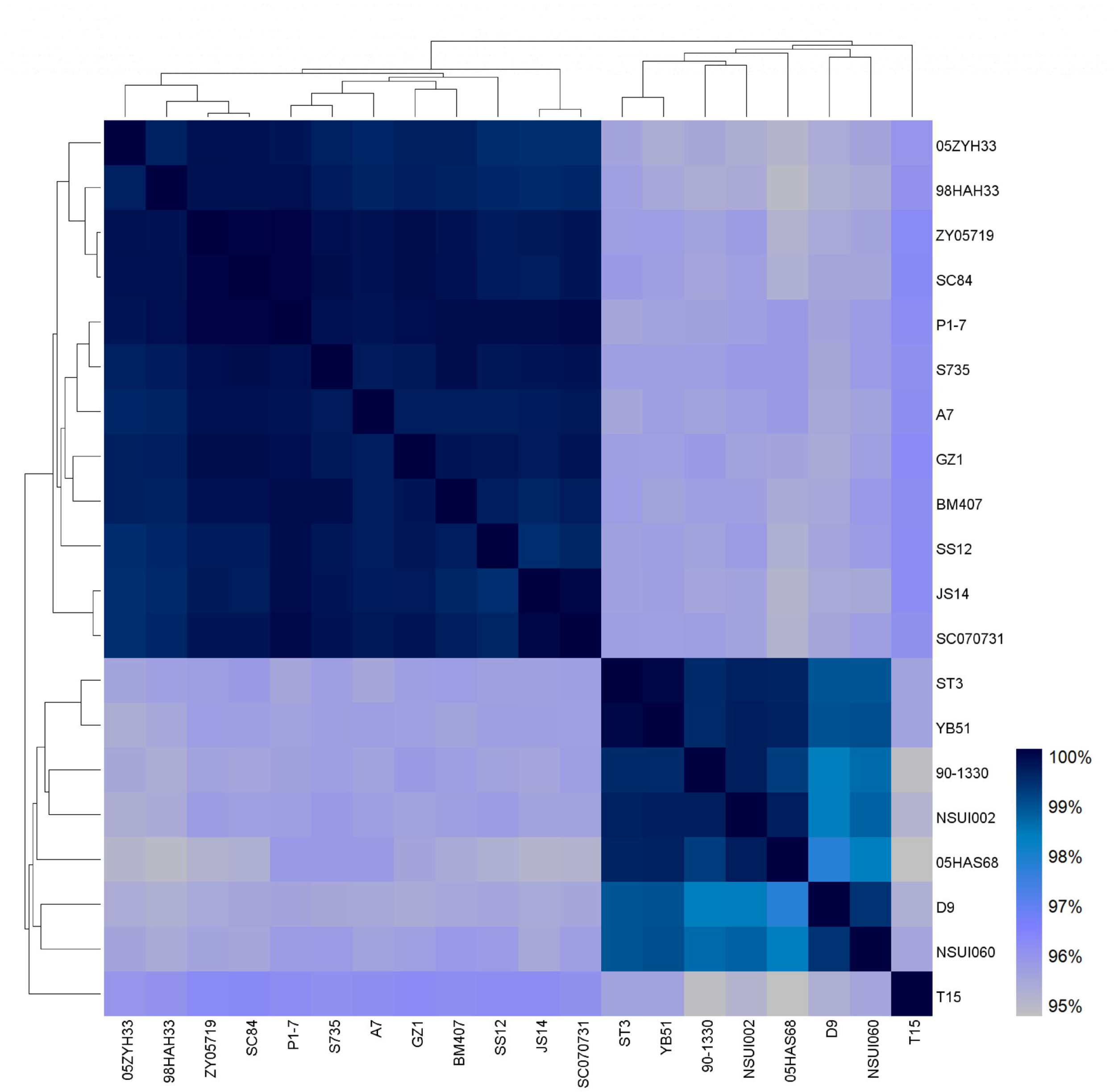
Heat map based on ANI values between every two genome sequences. The average nucleotide identity (ANI) were calculated using the web-based EZ BioCloud platform. Heatmaps based on ANI values were generated with HemI 1.0 (Heatmap Illustrator software, version 1.0).

### 3.5 SS3, SS7 and Clade 2 SS2 shared defense systems different from Clade 1 SS2

The defense elements, including RM system and CRISPR/Cas system, were detected in *S. suis* genomes. Both of the systems have a role in protecting bacteria against invading exogenous DNA, but the mechanism is different (Dupuis et al., 2013). The type I RM system uses HsdS, a single protein can respond to the methylation of target sequence, to determine the specificity of both restriction and methylation by the action of endonuclease HsdR and methyltransferase HsdM (Willemse and Schultsz, 2016). In *S. suis*, a total of three type I RM systems are detected in *S. suis* (Figure 6A). The Type A and Type B RM systems specifically appear in the Clade 1 SS2 strain, but not in Clade 2 strains. Type C RM system is present and conserved in strains of both clades. However, Clade 2 strains have an inserted *fic* gene between the *hsd* genes, which is different from Clade 1 strains. Another defense element, CRISPR/cas system, provides acquired immunity in prokaryotic organisms. It integrates short sequences of invading exogenous DNA between CRISPR repeats, and cleaves reinvading foreign DNA when recognize same sequences (Wiedenheft et al., 2012). CRISPR analysis result showed that the CRISPR components are absent in Clade 1 strains but present in all Clade2 strains. The CRISPR repeat in Clade 2 strains is 36 bp in length (GTTTTACTGTTACTTAAATCTTGAGAGTACAAA AAC), but SS7 strain D9 has an additional variant form with an additional TTA at the end of the repeat (Figure 6B and C). Above data is in accordance with a previous report (Okura et al., 2017), that SS3 and SS7 strains shared defense elements with Clade 2 SS2 strains, which is different from Clade 1 SS2 strains.

**Figure 6.**
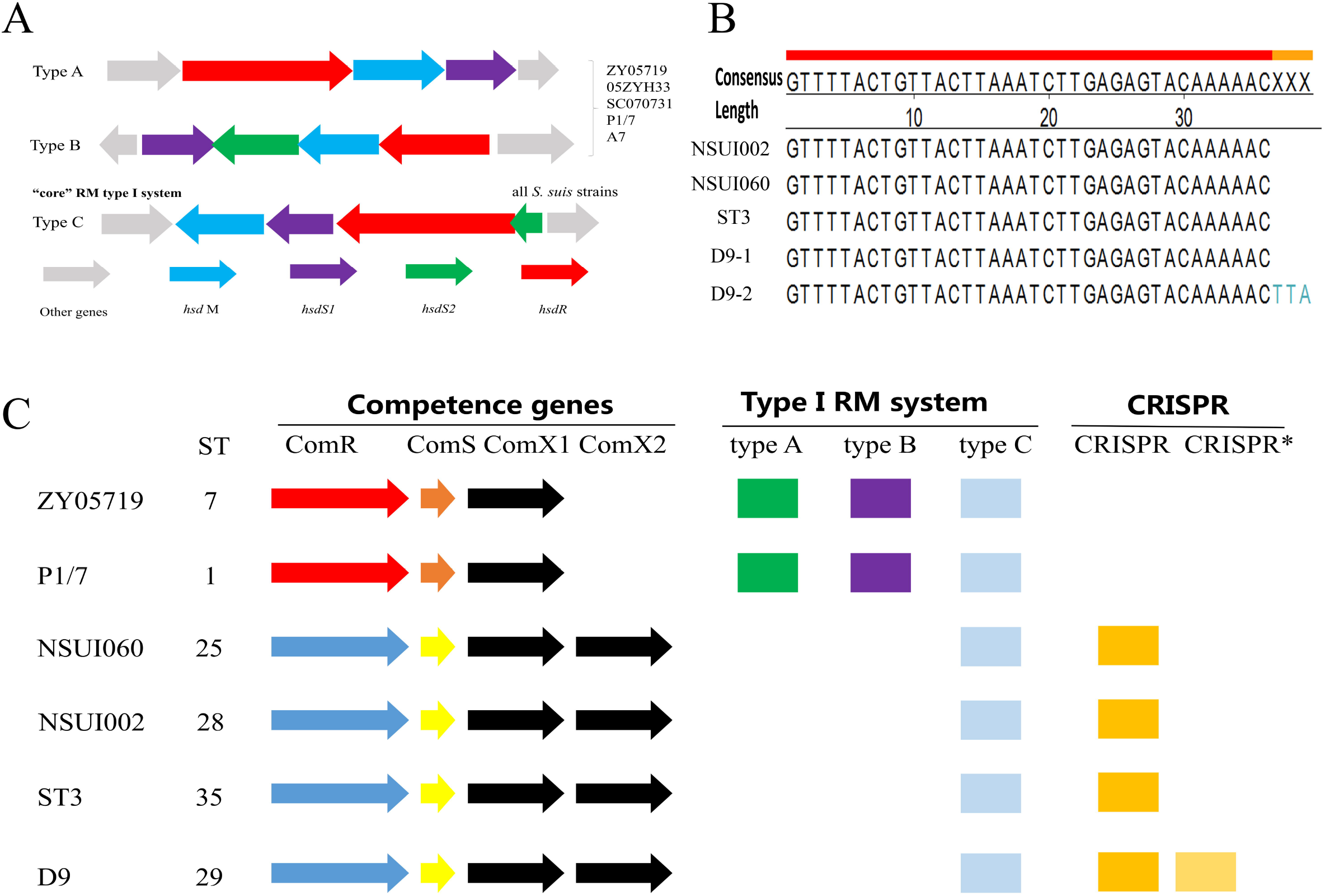
The structures of defense systems in *S. suis* isolates. **(A)** Three types of type I RM systems are found in *S. suis*. Type A and B RM systems only present in Clade 1 *S. suis* strains, and Type C was common RM system appearing in all of *S. suis* strains (Both Clade 1 and Clade 2). **(B)** Sequences of CRISPR repeats in representative *S. suis* strains. **(C)** The structures of competence and CRISPR elements in *S. suis*. Only Clade 2 strains harbors CRISPRs. Clade 1 and Clade 2 strains have different sequences in ComRS competence systems. The ComX regulator are conserved in *S. sui* strains, but Clade 2 strains have one more copy of ComX.

### 3.6 SS3, SS7 and Clade 2 SS2 shared a competence system different from Clade 1 SS2

*S. suis* is a bacterium with natural transformation ability, which depends on the ComRS competence systems. Natural transformation contributes to the horizontal gene transfer in *S. suis* and increase the diversity of *S. suis* genomes, which confers a unique advantage for capsule switch. Previously, we have identified three types of ComRS systems with specific competence pheromone in *S. suis* (Zhu et al., 2019). BLAST search results suggested that all Clade 1 isolates harbors Type A ComRS system, while all Clade 2 strains harbors Type B ComRS system (Figure 6C). We further detected the distribution of ComRS systems in field strains used in virulence genotyping (Figure 1). Results showed that all field strains that classified into cluster I in virulence genotyping have Type A ComRS system, whereas all strains classified into cluster II, including SS3 and SS7 isolates, harbor Type B ComRS system (data not shown). To experimentally test the transformation efficiencies of strains stimulated with noncognate synthetic competence pheromones, we randomly selected two cluster II SS2 strains (ZJJX0908005 and ZJ92091101), two cluster II SS3 strains (128-1-2 and 129-1-3), two cluster II SS7 strains (SH59 and SH04815) and two representative cluster I SS2 strains (ZY05719 and P1/7). Type A XIP exclusively induces competence in cluster I SS2 strains, and Type B XIP induces competence in cluster II SS2 strains, SS3 strains and SS7 strains (Figure 7). Although belonging to serotype 2, cluster II SS2 shared a same bacterial communication language with SS3 and SS7, but could not respond to the pheromone from cluster I SS2 strains, which is consistent with phylogenetic analyses results.

**Figure 7.**
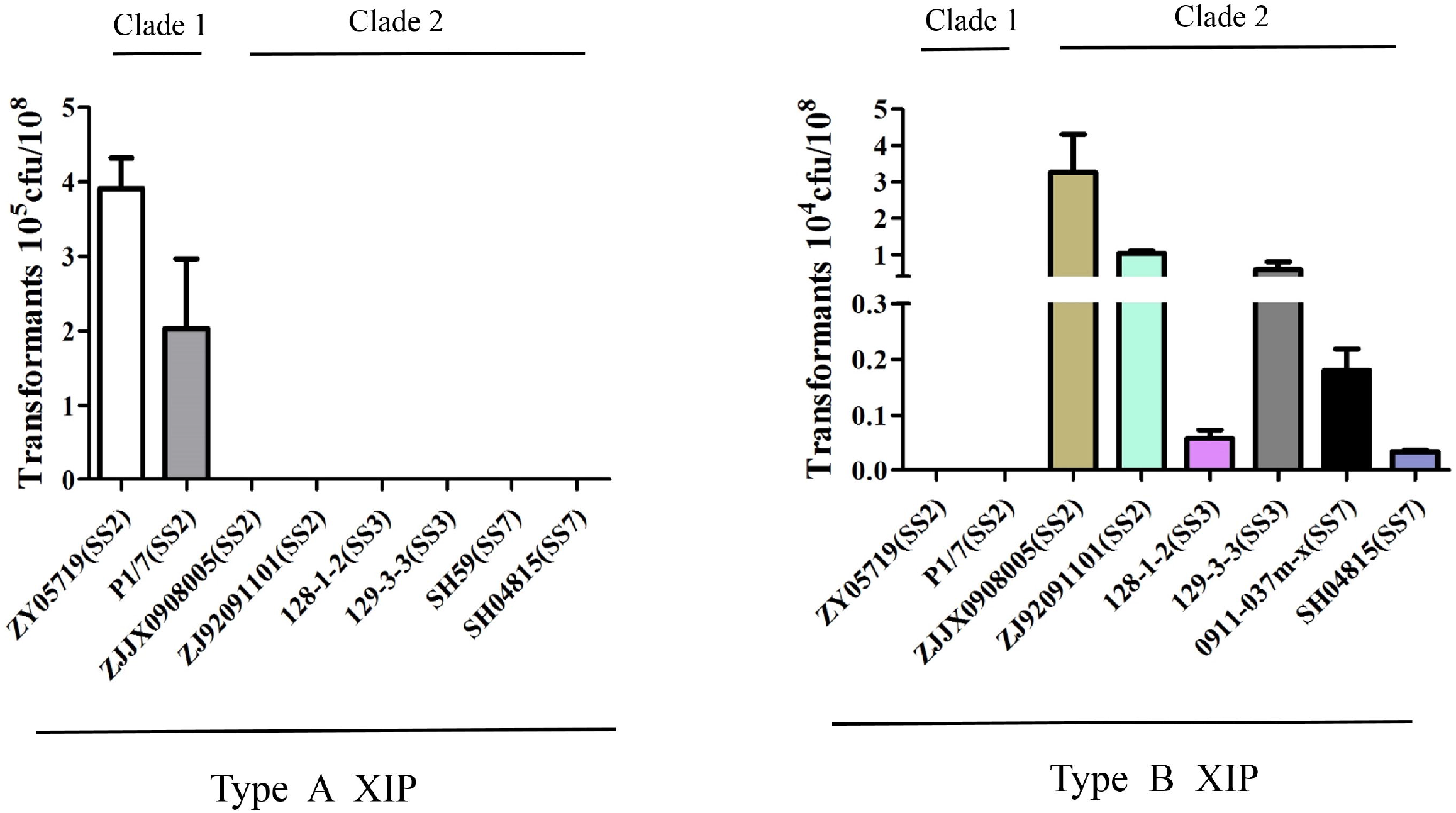
*S. suis* transformation induced by two different XIPs. Peptide XIP induces *S. suis* competence to uptake exogenous DNA. The competence effieiency can be assessed by positive transformants grow on THB (*spc:cm*+). Type A XIP is only able to induce transformation in Clade 1 SS2 strains. Type B XIP induces transformation in SS2, SS3, SS7 strains belonging to Clade 2.

### 3.7 Analysis of *cps* locus provided evidence for capsule switch between SS3 and Clade 2 SS2

To precisely characterize the capsular switch between SS2, SS3 and SS7, we compared the *cps* genes locus of the representative *S.suis* strains. Although SS7 strain D9 is closely related to ST25 SS2 strain NSUI060, BLAST search on the D9 genome showed that the *cps* locus is disrupted due to genomic rearrangement, and the *cps* genes are separated and translocated to different genomic segments (data not shown). Therefore, here we only compared representative SS2 and SS3 strains, including SS3 strain ST3, Clade 1 SS2 strain ZY05719 and Clade 2 strain NSUI002 (Figure 8). Sequence alignment of *cps* locus and its flanking regions of these strains revealed that the 7.2 kb-long upstream flanking sequence (*tetR* to *yaaA* and *cpsABCD*) and the 12.6 kb-long downstream flanking sequence (*aroA* to *asnS*) are extremely conserved among *S. suis* isolates. This suggested that the potential capsule switch event may occur through homologous recombination in these regions probably independently of the overall genetic background. Except for a translocated insertion of transposon element (red arrow), the SS2 *cps* locus of NSUI002 is almost identical to that of ZY05719, and differs from SS3 *cps* locus. However, in the ending region of SS2 *cps* locus, genes *cps2T, cps2U* and *cps2V* are absent from the *cps* locus of NSUI002. Instead, NSUI002 has a fragment shared high identity with sequence from SS3 *cps* locus containing *cps3O, cps3P* and *tnp3-4* (red line box). It is worth noting this unique region has no homologous sequence in the genome of ZY05719. Thus, this trait in the *cps* locus of NSUI002 strongly supports the hypothesis of capsular switch between SS2 and SS3, and favors a possibility that the potential capsular switch between SS3 and SS2 might result from a recombinational crossover point located ahead of *cps3O*. In summary, we find detailed evidence for capsule switch within *S.suis*, at least and as expected, from the analysis of c*ps* locus in CC28 SS2 and SS3 strains.

**Figure 8.**
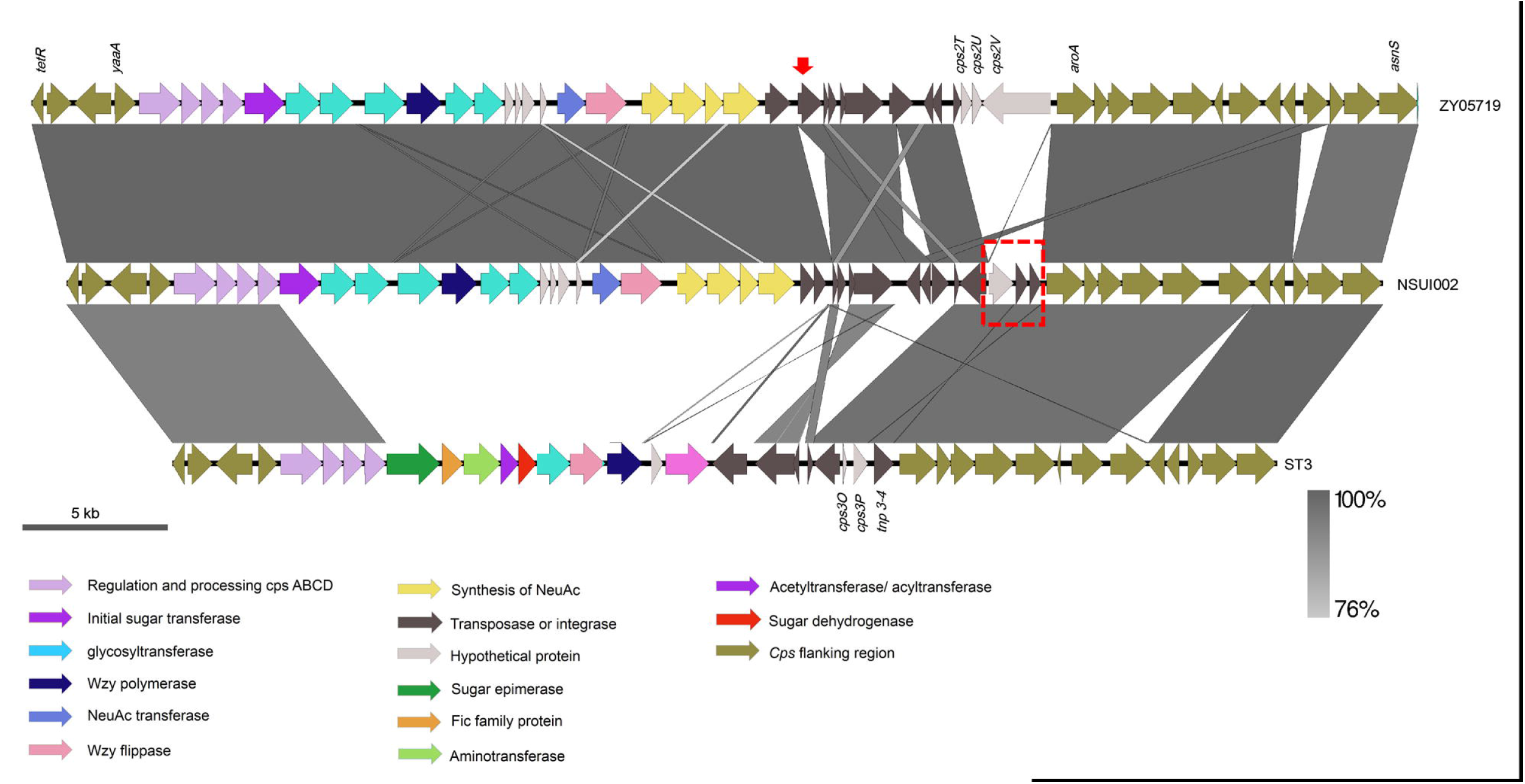
Schematic representations of *cps* locus in SS2 and SS3 strains. The *cps* locus of Clade 2 SS2 strain NSUI002, Clade 2 SS3 strain ST3 and Clade 1 SS2 strain ZY05719 were compared. The flanking regions of *cps* locus reveals high similarity. Red arrow indicates a translocation event. Red line box shows a region uniquely present in the *cps* locus of both ST3 and NSUI002.

## 4. Discussion

Serotype 2 is the most prevalent serotype of *S. suis* worldwide. Among the major evolutionary lineages revealed by genetics analyses, ST1 and 7 (CC1) SS2 are generally associated with diseases (Goyette-Desjardins et al., 2014). However, ST25 (CC29) and ST28 (CC28), accounting for larger proportions of SS2 strains in North American, present less virulence potentials in animal model (Athey et al., 2015; Fittipaldi et al., 2011). In accordance, ST28 strains in China are also regard as representive avirulent strains (Guo et al., 2020; Ma et al., 2020; Wang et al., 2017). Those reports suggested that CC28 and CC29 SS2, which are phylogenetically distinct from CC1 SS2, are the pool of strains with lower virulence levels. In this study, we report that capsule switch exists in *S. suis* population, notably in CC28 and CC29 between SS2, SS3 and SS7. This finding may explain the genetic and phenotypic differences between CC1 SS2 and CC28/29 SS2, and indicate a possibility that CC28/29 SS2 was derived from an ancestor unrelated with CC1 SS2 through capsule switch, which has often been overshadowed by simple serotyping.

Capsule switch has been identified in other extracellular pathogens harboring polysaccharide capsule and natural competence, such as *Neisseria meningitidis (N, meningitidis), S. agalactiae* and *S. pneumoniae*. Among which, the capsule switch of *S. pneumoniae* is the most representative and well-studied. Pneumococcal capsule switch can be achieved through gradual evolution with a combination of minor mutation, deletion and recombination in *cps* locus. For example, pneumococcal serotype 6A, 6B, 6C and 6D have near identity of *cps* locus, which only differ in *wciP* gene (Song et al., 2011). A similar case in *S. suis* is the capsule switch between SS1, SS2 and SS1/2 in CC1. Serotype 1/2 cannot be differentiated from serotype 1 and 2 by serum antigenicity (Perch et al., 1983), and genetic analysis of the *cps* locus demonstrated serotypes 1, 1/2 and 2 share the high genetic identities (Okura et al., 2013). On the other hand, capsule switch can occur through the exchange of large genomic fragment containing full *cps* locus (Bellais et al., 2012; Wyres et al., 2013), which is a suitable model to be applied in the capsule switch between SS 2, 3, 7 strains in CC28 and 29. Given the fact that *cps* locus is high diverse and variable between different *S. suis* serotypes (Okura et al., 2013), it is not logical to keep *cps* locus conserved or even identical when genetic backbone is subject to high degree of genetic rearrangement. Therefore, our data strongly supports a hypothesis that serotype 2, 3, 7 capsule switch results from the exchange of large *cps* locus, leads to similar genetic backbones sharing common competence systems and defensive systems.

Importantly, the upstream and downstream flanking sequences of *cps* locus are almost identical between different *S. suis* serotypes (Okura et al., 2013), providing potential recombination sites for capsule switch. Furthermore, we studied the sequence of *cps* locus and flanking region between a CC1 SS2 ZY05719, CC28 SS2 strain NSUI002 and SS3 strain ST3 in detail. A genetic fragment was found to be conserved in the *cps* locus of CC28 SS3and SS2, but absent at that location in CC1 SS2, which suggests a potential recombination event occurred between CC28 SS2 and SS3. However, based on current information, we cannot determine the temporal relationship between strains, namely whether one is derived from another, or there is a common ancestor. In addition, although ST27 (a ST harboring both SS2 and SS3), ST28 (including NSUI002, SS2) and ST35 (including YB51 and ST3, SS3) are phylogenetically related and clustered together in CC28, ST35 SS2 or ST28 SS3 have not yet been observed, indicating that additional events occurred after capsule switch in this common evolutionary lineage. More collected isolates and sequenced genomes in the future would be helpful to address those issues.

In fact, the core of capsule switch is to increase diversity in the population and enhance fitness in certain environments, such as increasing antibiotic resistance or escaping herd immunity. Accumulated studies have demonstrated that capsule switch in *S. pneumonia* can be attributed to the selection pressure from the use of vaccines targeting capsule (Martins et al., 2010). Similarly, the change of CPS composition caused by *cpsG* mutation in *S. iniae* and capsule switch (from type III to IV) in CC17 *S. agalactiae* contributes to vaccine escape (Bellais et al., 2012; Heath et al., 2016). Besides that, capsule switch is also involved in bacterial pathogenicity. Pneumococcal capsule switch from 6A to 6C promotes the resistance to complement system and presents enhanced virulence for respiratory tract infection (Sabharwal et al., 2014). Artificial capsule switch in *S. pneumonia* is able to alter virulence and infection outcomes in a mouse model (Kelly et al., 1994; Trzcinski et al., 2003). Although the role of capsule switch in *S. suis* is not clear, it is possible that the replacement of *cps* locus would cause phenotypic changes due to the alteration of bacterial surface architecture. For instance, *NeuB*, an enzyme existing in serotype 1/2, 2 and 14, is essential for sialic acid biosynthesis (Feng et al., 2012). Therefore, acquiring NeuB may increase bacterial resistance to complement system and phagocytosis (Feng et al., 2012; Uchiyama et al., 2019). In fact, a recent study highlighted that experimentally switching capsule type 2 to 3 in *S. suis* leads to defective whole blood survival and bacterial virulence (Okura et al., 2020). More studies in the future are needed to clarify the phenotypic features of different *S. suis* capsules, and what benefits bacteria may obtain from capsule switch.

## Acknowledgements

This work was funded by the Natural Science Foundation of China (31972650), Shanghai Agriculture Applied Technology Development Program (S0201700386), the Key Project of Independent Innovation of the Fundamental Research Fund for the Central Universities of Nanjing Agricultural University (KJQN201932) and the Priority Academic Program Development of Jiangsu Higher Education Institutions (PAPD). We would like to thank Dr. Qiang Li for his help in bioinformatic analyses.

## Data availability statement

The data that support the findings of this study are available from public database GENBANK, and the accession number is listed in Table. S2

## Ethical approval

No animal experiments were performed thus ethical statement is not applicable in this study.

