## Supplementary tables for "Comparative genomic analyses provide clues to capsule switch in *Streptococcus suis*"

**Table S1** The results of MLST for SS2, SS3 and SS7 strains

| Strain | Serotype | ST |
| --- | --- | --- |
| P1/7 | 2 | 1 |
| GZ1 | 2 | 1 |
| ZY05719 | 2 | 7 |
| SC84 | 2 | 7 |
| 05HAS68 | 2 | 28 |
| 90-1330 | 2 | 28 |
| NSUI002 | 2 | 28 |
| NSUI060 | 2 | 28 |
| P4254 | 2 | 28 |
| LS091105 | 2 | 28 |
| HZ060601 | 2 | 28 |
| ZJJX0908005 | 2 | 28 |
| HN1004001 | 2 | 28 |
| ZJ92091101 | 2 | 28 |
| HN0104001 | 2 | 28 |
| 0911-065m-2 | 7 | 225 |
| 0911-065m-2 | 7 | 225 |
| ST3 | 3 | 35 |
| YB51 | 3 | 35 |
| 128-2-1 | 3 | 27 |
| 129-3-3 | 3 | 27 |
| 128-1-1 | 3 | 27 |
| ZJNB150 | 3 | 265 |
| HN147 | 3 | 117 |
| ZW43 | 3 | 117 |
| HN121 | 3 | 117 |
| ZW31 | 3 | 117 |
| SS13 | 3 | 117 |
| ZW29 | 3 | 117 |
| Hb1001 | 3 | 117 |
| D9 | 7 | 29 |
| SH59 | 7 | 225 |
| SH04815 | 7 | 225 |
| SH04805 | 7 | 225 |

1 Phylogenetic tree conducted by analyzing the concatenation sequences of MLST data in this table is shown in Figure S2.

**Table S2** Characteristics of various *S. suis* strains used for analysis in this study

| Strain | Serotype | Length(bp) | GC% | CDS | GenBank ID |
| --- | --- | --- | --- | --- | --- |
| BM407 | 2 | 2146229 | 41.1 | 1947 | NC_012926.1 |
| 05ZYH33 | 2 | 2096309 | 41.1 | 2186 | NC_009442.1 |
| 98HAH33 | 2 | 2095698 | 41.1 | 2185 | CP000408.1 |
| GZ1 | 2 | 2038034 | 41.4 | 1872 | NC_017617.1 |
| SC84 | 2 | 2095898 | 41.1 | 1973 | NC_012924.1 |
| P1/7 | 2 | 2007491 | 41.3 | 1898 | NC_012925.1 |
| A7 | 2 | 2038409 | 41.2 | 1891 | NC_017622.1 |
| S735 | 2 | 1980887 | 41.4 | 1840 | NC_018526.1 |
| SC070731 | 2 | 2138568 | 41.2 | 2003 | NC_020526.1 |
| ZY05719 | 2 | 2094898 | 41.1 | 1926 | NZ_CP007497.1 |
| T15 | 2 | 2240234 | 41.0 | 2100 | NC_022665.1 |
| 05HAS68 | 2 | 2188363 | 41.2 | 2065 | NZ_CP002007.1 |
| 90-1330 | 2 | 2146151 | 41.0 | 2064 | NZ_CP012731.1 |
| NSUI002 | 2 | 2255345 | 41.1 | 2221 | CP011419 |
| [NSUI060](http://www.ncbi.nlm.nih.gov/genome/199?genome_assembly_id=265872) | 2 | 2302206 | 41.1 | 2185 | NZ_CP012911.1 |
| SS12 | 1/2 | 2096866 | 41.2 | 1989 | NC_017619.1 |
| ST1 | 1 | 2034321 | 41.4 | 1869 | NC_017950.1 |
| ST3 | 3 | 2028815 | 41.2 | 1952 | CP002633.1 |
| YB51 | 3 | 2043228 | 41.2 | 2012 | NC_022516.1 |
| 6407 | 4 | 2292360 | 41.0 | 2127 | NZ_CP008921.1 |
| D9 | 7 | 2277656 | 41.0 | 1966 | NC_017620.1 |
| D12 | 9 | 2183059 | 41.3 | 2008 | NC_017621.1 |
| DN13 | 9 | 2142184 | 41.3 | 2021 | NZ_CP015557.1 |
| JS14 | 14 | 2137435 | 41.2 | 1979 | NC_017618.1 |
| TL13 | 16 | 2038146 | 41.3 | 1874 | NC_021213.1 |
