## Supplementary figures and images for "Comparative genomic analyses provide clues to capsule switch in *Streptococcus suis*"

### Supplementary Figure 1

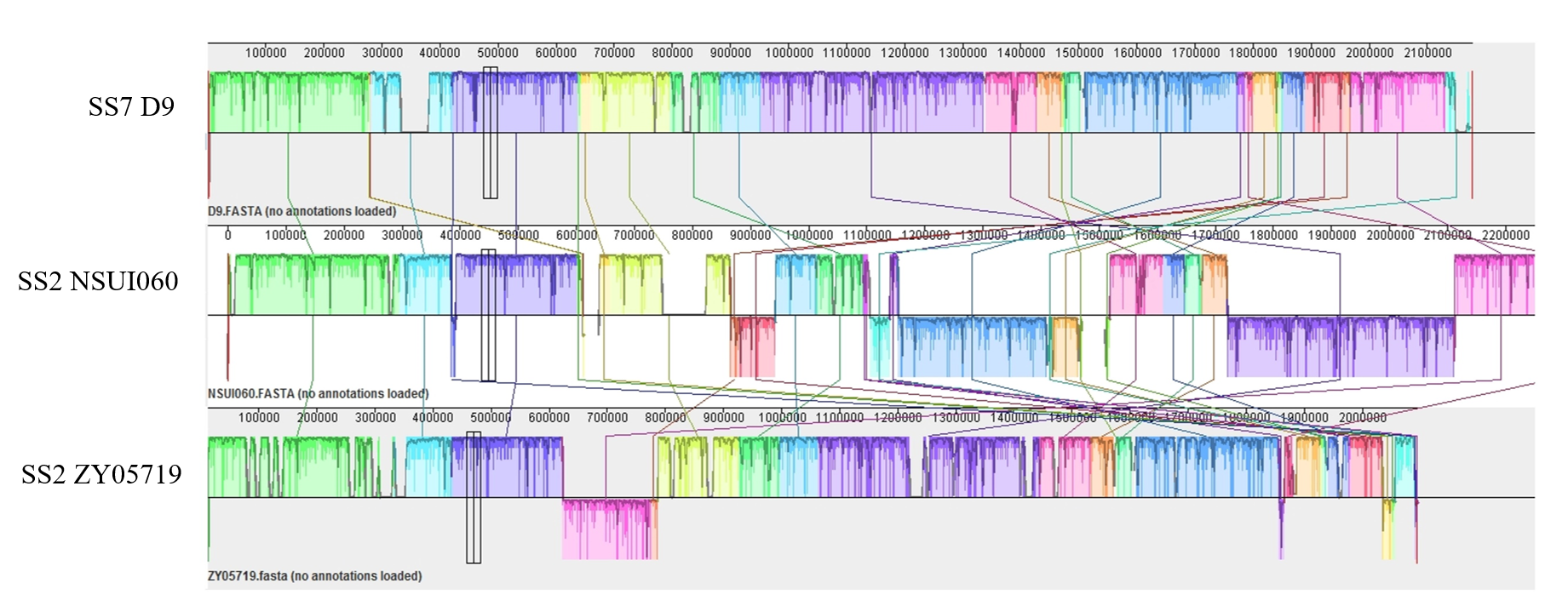

### Supplementary Figure 2

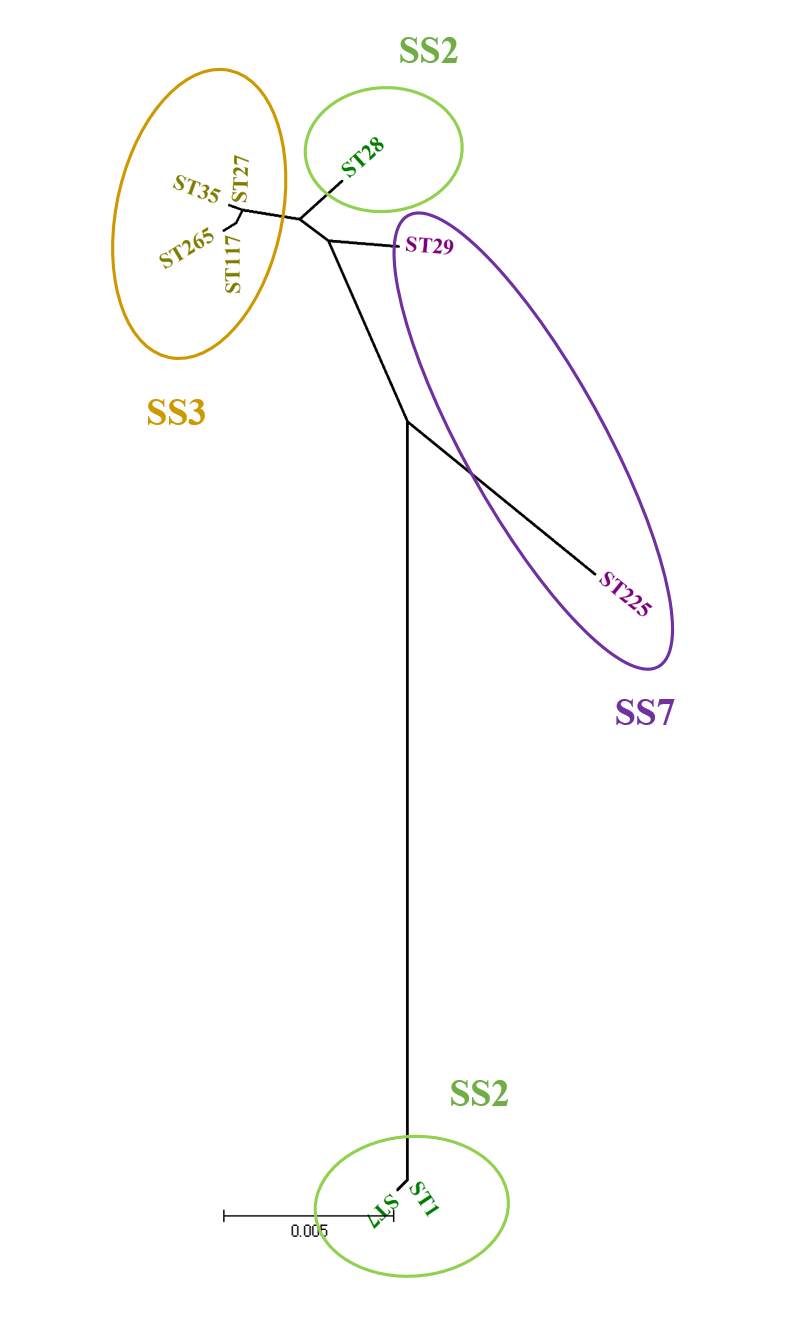
